# Maturation of Human Intestinal Epithelial Cell Layers Fortifies the Apical Surface against *Salmonella* Attack

**DOI:** 10.1101/2024.07.11.603014

**Authors:** Jorik M. van Rijn, Ana C. C. Lopes, Merve Ceylan, Jens Eriksson, Alexandra Bergholtz, Angelika Ntokaki, Rebekkah Hammar, Magnus Sundbom, Martin Skogar, Wilhelm Graf, Dominic-Luc Webb, Per M. Hellström, Per Artursson, Thaher Pelaseyed, Maria Letizia Di Martino, Mikael E. Sellin

## Abstract

The active invasion of intestinal epithelial cells (IECs) represents a key event in the infection cycle of many gut pathogens. Studies of how *Salmonella enterica* Typhimurium (*S*.Tm) bacteria enter transformed cell lines have shaped the paradigm for swift type-three-secretion-system-1 (TTSS-1)-driven IEC invasion, fueled by expansive membrane ruffles. However, comparative studies suggest that non-transformed IECs in the intact gut context comprise a much more challenging target for the attack. The molecular and cellular features that explain these discrepancies remain undefined. By live-cell imaging in human enteroid-and colonoid-derived IEC layers, we demonstrate that the maturation state of gut epithelia dramatically impacts permissiveness to *S*.Tm invasion. IEC layers kept under immature progenitor-cell-promoting conditions are permissive to the bacterial invasion, whereas maturation towards an enterocyte/colonocyte fate reduces the frequency of *S*.Tm-induced epithelial entry structures, and lowers the invasion efficiency by up to tenfold. This phenotypic shift during IEC maturation couples to an altered expression of actin regulatory proteins implicated in the invasion process, and an increased dependence on the *S.*Tm TTSS-1 effector SipA for successful entry. In addition, IEC maturation involves upregulation of cell surface mucins, e.g. MUC13, and shifts in glycocalyx composition, as revealed by multiple lectin stainings. Enzymatic treatment of the apical surface with the StcE mucinase converts maturing IEC layers back to the *S*.Tm-invasion-permissive state of their immature counterparts. Taken together, these results showcase how the maturation state of human IECs dictates the susceptibility to invasion by a prototype enterobacterium.

## Introduction

Enteropathogenic microorganisms can attack and often invade into intestinal epithelial cells (IECs), a feature distinguishing these agents from the commensal gut microflora. IEC invasion is a critical step in the pathogenesis of several enteropathogenic bacteria, including *Salmonella*, *Shigella*, enteroinvasive *Escherichia coli*, and *Listeria* strains. The capacity to invade IECs enables these bacteria to access additional host niches for replication and long-term colonization, but also triggers an acute inflammatory response in the infected mucosa (1–3). Due to the pivotal role of this infection step, the underlying mechanisms have been under intense investigation during the last decades.

To decipher the molecular interactions between invasive bacteria and host cells, researchers have turned to experiments in cultured cell lines, e.g. the human cervical carcinoma line HeLa, human colorectal carcinoma lines such as Caco-2 and HT-29, or canine kidney MDCK cells (2). These have a number of advantages, e.g. ease of handling, powerful tools for perturbation of host cell gene expression, and the possibility to grow the cells flat on glass surfaces compatible with high-resolution live-cell bioimaging. Studies of *Salmonella enterica* Serovar Typhimurium (*S*.Tm) and related enterobacteria in such models have resulted in a detailed understanding of bacterium-triggered IEC invasion (1–3). *S*.Tm reaches the epithelial surface by flagellar motility, and deploys adhesins in combination with a type-three-secretion-system (TTSS-1) to dock to the host cell surface (4–10). The TTSS-1 subsequently delivers multiple effectors into the host cell to drive activation of Rho-GTPases, Arf-GTPases and formins, fueling actin polymerization and bundling (11–19). The TTSS-1 effectors SopB, SopE, and SopE2 appear particularly influential for *S*.Tm invasion of epithelial cell lines (10, 20). Their combined actions cause the near-instantaneous emergence of expansive membrane ruffles for bacterial uptake. Collateral ruffling also sparks cooperative uptake of nearby bacteria (20–23).

A conceptual framework for how *S*.Tm and related enterobacteria invade IECs thus exists. However, since the underlying data rely on cell line experiments, it remains poorly understood how the features and context of primary IECs affect the invasion process. Tumor-derived cell lines are fast-growing, exhibit a high degree of dynamics, and when grown flat on glass or plastic surfaces lack both the confluent polarized cell arrangement, and the densely microvilliated and heavily glycosylated apical surface structures characteristic of enterocytes and colonocytes (24, 25). Notably, in recent comparative work, stark differences were indeed observed between how *S*.Tm invades cultured cell lines versus absorptive IECs *in vivo* in mice (10). In the murine gut, *S*.Tm invasion of IECs relies strongly on the TTSS-1 effector SipA with a less pronounced contribution of SopB, SopE, and SopE2, and elicits markedly smaller, more transient entry structures than the prototypical ruffles fostering cooperative invasion in cell lines (10). Moreover, the giant adhesin SiiE appears redundant for *S*.Tm invasion of flat-growing cell lines such as HeLa. In contrast, SiiE is of key importance for the bacterium to invade polarized IEC layers and the murine gut epithelium *in vivo*, by mediating interactions with cell-surface-bound mucins (5, 7, 10). These observations suggest that while commonly used cell line models are highly permissive to *S*.Tm invasion and allow for rapid ruffle-mediated entry, *S*.Tm invasion of primary absorptive IECs is significantly more challenging (10, 26). The basis for these differences, and whether the conclusions from mice extrapolate fully to human intestinal epithelia, remain unclear.

Organoids grown from either embryonic, induced, or adult stem cells are currently revolutionizing infection biology (26–35). “Enteroids” and “colonoids” derive from adult stem cells extracted from small intestinal, or colonic, intestinal epithelial crypts. They develop into pure, non-transformed, intestinal epithelial cultures that can be maintained from both mouse and human in a variety of differentiation states and geometries (26, 32, 36–40). Hence, these models offer attractive experimental possibilities for the study of pathogen interactions with a near-native epithelium (41–43). We have recently developed technology to grow 2D intestinal epithelial cell layers from enteroids inside a custom-designed “Apical Imaging Chamber”. This allows tracing of enteropathogen attack on the epithelial surface at high temporal and spatial resolution (26). Here, we combined this approach with protocols to culture human enteroid-and colonoid-derived IEC layers under defined differentiation conditions. Our findings reveal that immature human IECs, in analogy to tumor-derived epithelial cell lines, remain relatively permissive to *S*.Tm invasion. By sharp contrast, IEC maturation towards an enterocyte or colonocyte phenotype causes shifts in the proteomes, apical surface structure, and glycocalyx composition that together fortify the epithelium against the bacterial attack.

## Results

### A Panel of Human Intestinal Epithelial Cell Layers with Distinct Maturation States, Compatible with Live-Cell Imaging at the Apical Surface

Building on the recently developed Apical Imaging Chamber (AIC (26)), we established a panel of enteroid-and colonoid-derived intestinal epithelial monolayers, compatible with live-cell imaging. First, human enteroids were expanded in Matrigel domes overlaid with Wnt3a/EGF/Noggin/R-spondin-1-containing organoid growth medium (OGM). The enteroids were extracted from the domes, fragmented, and the cell suspension seeded in OGM onto serially precoated alumina membranes mounted in AICs (or onto polyethylene terephthalate (PET) transwell inserts for some applications; see Methods). At day 3 of culture, the monolayer medium was exchanged for either i) fresh OGM, ii) medium deprived of Wnt3a, but still containing EGF, Noggin, and R-spondin (ENR), or iii) the ENR medium supplemented with RANK ligand and Tumor necrosis factor (ENRRT) (**Fig 1A**). We predicted that this would fuel intestinal epithelial cell (IEC) differentiation along unique trajectories. Indeed, at 7 days post-seeding, the three monolayer types exhibited distinct transcriptional profiles (Principal component analysis in **Fig S1**), compatible with an immature progenitor cell state (OGM), a more mature enterocyte state (ENR), or a state enriched in M-cell-associated transcripts (ENRRT), respectively (**Fig 1B**). By confocal fluorescence and scanning electron microscopy (SEM), OGM monolayers were found to feature an undulating sparsely microvilliated surface, whereas the ENR monolayeŕs surface was densely microvilliated (**Fig 1C**). The ENRRT monolayers displayed short aggregated surface protrusions or bundled microvilli, although the fully developed microfolds seen on M-cells *in vivo* were not observed (**Fig 1C**) (44). Under all three conditions, goblet cells appeared essentially absent, suggesting counterselection of secretory lineage cells in these settings (**Fig 1C**; see also proteomic results below). Finally, and further in line with distinct IEC states between conditions, the ENR monolayer displayed a higher nucleus to apical brush border distance than OGM or ENRRT, which was reproducible in monolayers generated from three distinct enteroid lines (**Fig 1D**). We conclude that human IEC monolayers can be established atop PET transwell inserts and in AIC chambers under defined conditions that yield either an immature progenitor cell state (OGM), a maturing enterocyte-like state (ENR), or monolayers partially enriched for an M-cell-like state (ENRRT).

**Figure 1.**
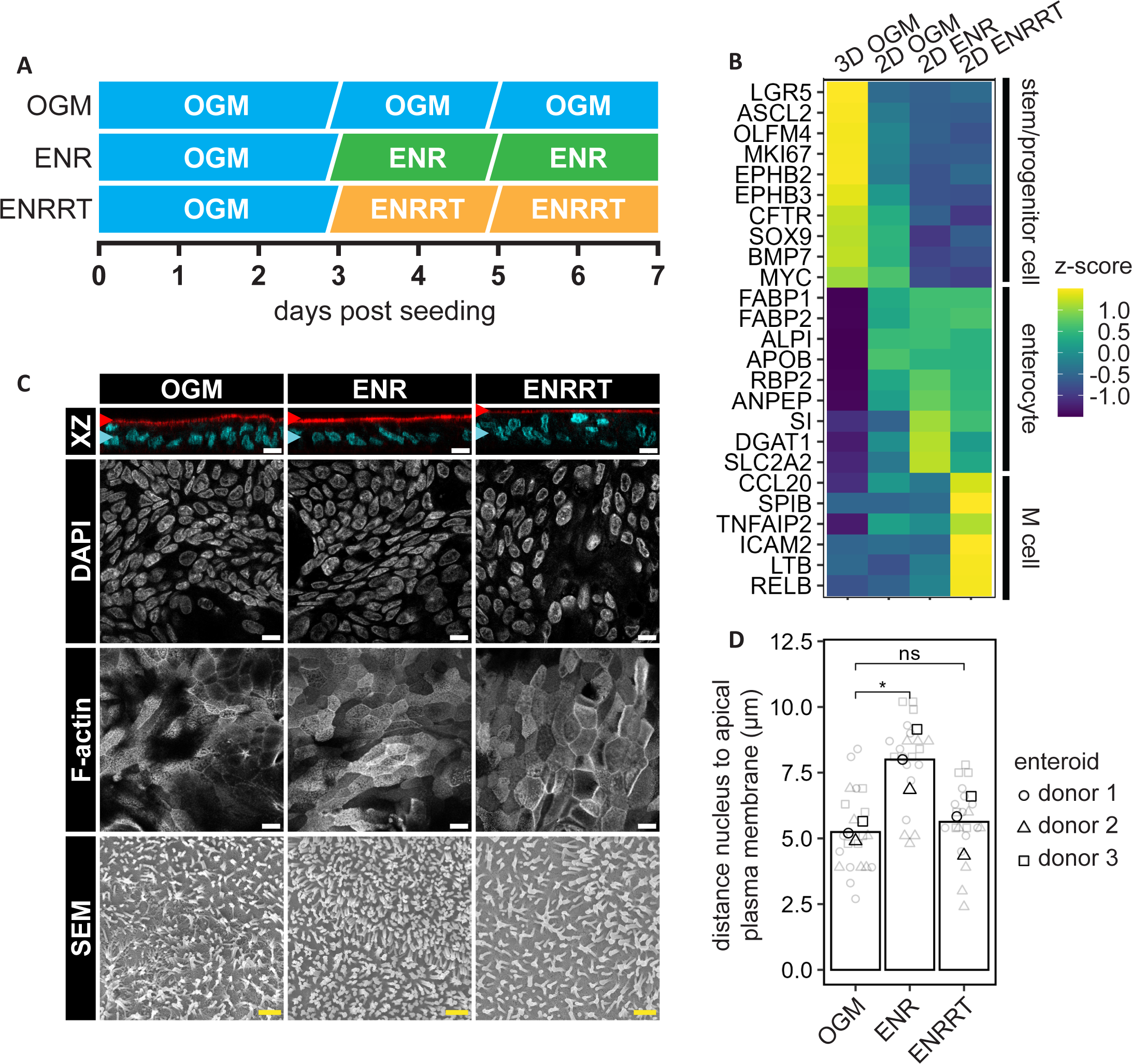
Directed differentiation of enteroid-derived intestinal epithelial cell layers results in distinct microscopic and transcriptional phenotypes. **(A)** Schematic of the differentiation strategy. 3D enteroids were expanded in OGM for 7-9 days, disrupted into single cells, and cultured as 2D monolayers on either coated PET transwell supports, or coated AICs. The monolayers were grown confluent in OGM for 3 days and, where applicable, differentiated in ENR or ENRRT for an additional 4 days. **(B)** Transcriptional profiles of 2D enteroid-derived monolayers. 3D enteroids in OGM included as control for a stem-cell rich phenotype. Data plotted as Z-scores. 2D monolayers grown in OGM exhibited a transitional state with weak stem cell marker expression (LGR5, ASCL2, OLFM4), reduced proliferation (MKI67), but retention of an immature phenotype (SOX9, BMP7, MYC) and only modest expression of enterocyte markers (FABP1/2, ALPI, APOB, RBP2, ANPEP). Incubation in ENR caused elevated enterocyte marker expression (particularly SI, DGAT1, SLC2A2), while ENRRT promoted M cell marker expression (CCL20, SPIB, TNFAIP2, ICAM2, LTB, RELB). **(C)** Representative micrographs of differentially cultured 2D enteroid-derived monolayers. Confocal microscopy (top, middle and XZ projection rows), following staining with DAPI (nuclei; teal in XZ projection) and phalloidin (F-actin; red in XZ projection) revealed a confluent monolayer for each condition with the majority of nuclei in the same plane, and with the apical brush border most distinctly evident for the ENR condition. Scanning electron microscopy (bottom row) showed a sparsely microvilliated surface for OGM-grown monolayers and denser microvilliation for the ENR condition. ENRRT-grown monolayers were characterized by short aggregated surface protrusions and/or bundled microvilli. White scale bars, 10µm. Yellow scale bars, 1µm. **(D)** Quantification of the nucleus to cell surface distance for differentially cultured 2D enteroid-derived monolayers from three donors. Monolayers cultured atop PET transwell supports were fixed and stained with DAPI (nuclei) and WGA (surface marker). In Z-stacks, the distance between nuclei centers and the cell surface was measured, which revealed apicobasal elongation of IECs in the ENR condition. For each donor culture, the mean distance was measured across three fields of view for each condition. Presented values are pooled from n=2 biological replicates within each condition for donors 2 and 3 and n=3 replicates within each condition for donor 1. Statistical analysis; unpaired, two-sided t-test with Holm-Bonferroni post-hoc using the OGM condition as reference. ns p-value = 0.676, *p-value = 0.0256.

### Human Intestinal Epithelial Cell Differentiation towards a Maturing Enterocyte or Colonocyte state Reduces Permissiveness to *Salmonella* Invasion

To address how the IEC state affects permissiveness to invasive bacteria, we next performed live-cell imaging in AICs on a custom-built microscope (26). The culture medium atop enteroid monolayers was replaced for DMEM/F12, and 10^6^ wild-type *Salmonella enterica* Typhimurium SL1344 (*S*.Tm *wt*) were inoculated in the apical AIC compartment, corresponding to a multiplicity of infection (MOI) of ∼3. Differential interference contrast (DIC) imaging was performed for 60min post-infection (p.i.) with the focal plane at the apical epithelial surface. This revealed a gradual accumulation of *S*.Tm atop the monolayers and exemplified the frequent lingering of the bacteria on the IEC surface (**Fig 2A-B**; Tracking in cyan), also noted in our recent study (26). From ∼10-20min p.i. onwards, examples of *S*.Tm invasion events became evident, signified by the emergence of entry structures of varying morphology on the apical IEC surface and the sinking in of the bacterium into said surface and out of the focal plane (**Fig 2A-B**). Strikingly, the frequency of *S*.Tm invasion events differed dramatically between the three monolayers differentiation states. In maturing ENR (enterocyte-like) monolayers, only ≤10% of surface-attached *S*.Tm invaded within the 60min imaging window (**Fig 2B-C**). By sharp contrast, invasion events were much more prevalent in the immature OGM monolayers (∼50% of surface-attached *S*.Tm; **Fig 2B-C**). M-cells *in vivo* are known for their capacity to take up luminal particles and microbes (44). In line with this, ENRRT monolayers were also highly permissive to *S*.Tm invasion (∼70% of all surface-attached bacteria) (**Fig 2B-C**). Hence, maintaining human IEC monolayers in an immature progenitor state, or exposing them to a medium that promotes differentiation along an M-cell trajectory favors *S*.Tm invasion, whereas maturation towards an enterocyte phenotype reduces permissiveness to invasion. These differences held true for quantification of triplicate OGM, ENR, and ENRRT monolayer infection experiments using the same enteroid line (**Fig 2C**), and were also verified using a separately established line (**Fig S2A**). To examine if these findings generalized across gut segments, we next established human colonoids and generated colonoid-derived OGM and ENR monolayers for imaging. Again, the OGM monolayers were markedly more permissive to *S*.Tm invasion than the ENR monolayers (∼70% vs ∼10% of surface-located *S*.Tm bacteria successfully invading by 60min p.i.) (**Fig 2B-C**; verified also for an additional colonoid line, **Fig S2B**).

**Figure 2.**
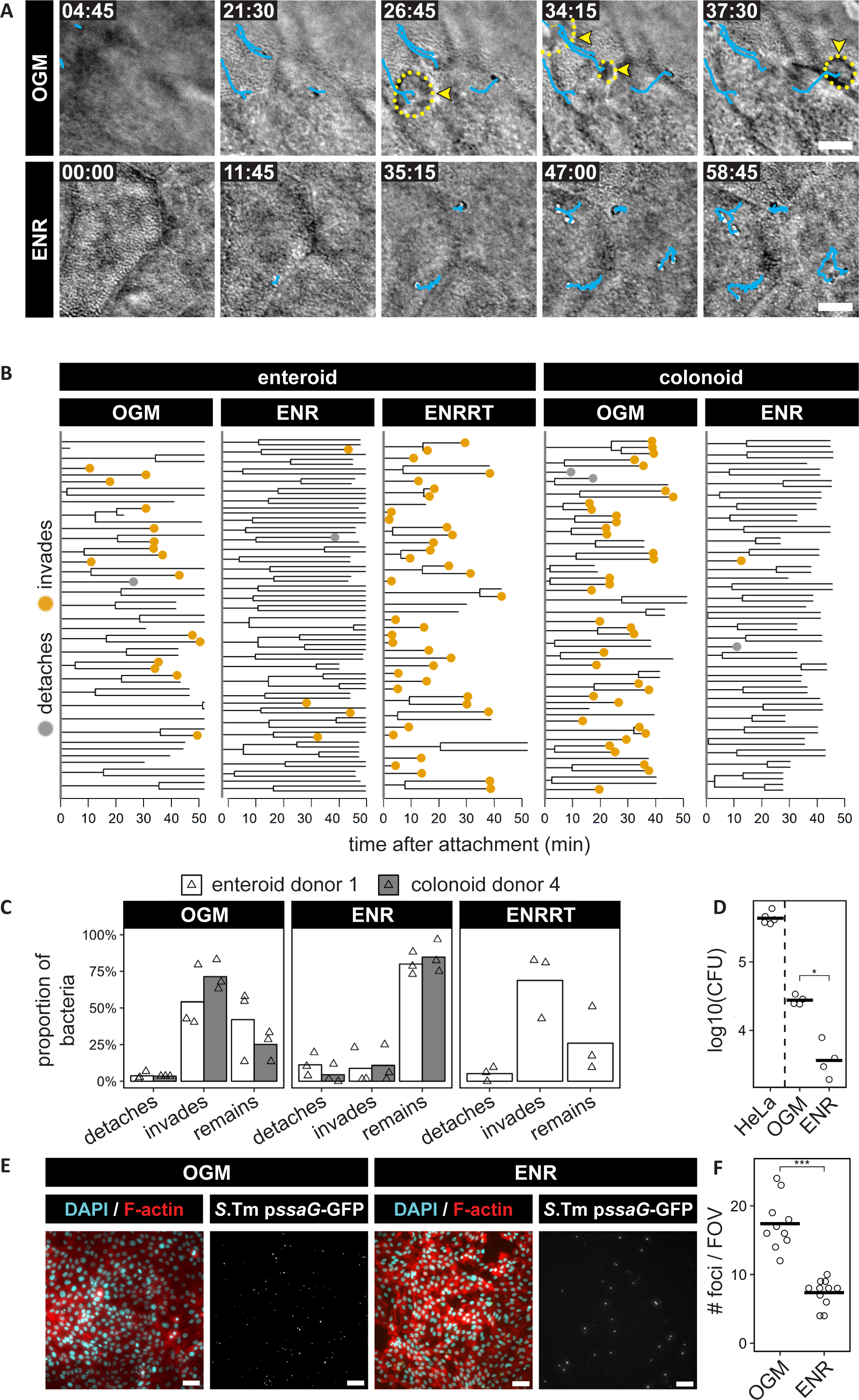
Maturation of human intestinal epithelial cell layers towards an enterocyte/colonocyte phenotype reduces permissiveness to *Salmonella* invasion. **(A-C)** Time-lapse imaging of enteroid-and colonoid-derived IEC monolayers grown in OGM, ENR or ENRRT within AICs and infected with *S*.Tm *wt* and MOI ∼3 for 1h. **(A)** Representative time-lapse DIC image series of *S*.Tm at the apical epithelial surface of OGM-(top row) and ENR-grown (bottom row) enteroid-derived monolayers. Cyan lines represent tracking of bacterial movements atop the monolayer. *S*.Tm invasion events indicated by yellow arrowheads and the entry structure areas outlined by dotted circles. Time in min:sec; scale bars, 10µm. **(B-C)** Quantification of the fates of individual *S*.Tm after initial attachment to the epithelial surface. In **(B)** the behavior of individual surface-bound *S*.Tm in a representative time-lapse movie from each condition is visualized as dendrograms. The dendrograms indicate the timing of bacterial division (track split), detachment (grey dot), invasion (yellow dot), and remaining bacteria at the end of the movie or loss of focus (bare branch). In **(C)** the proportion of *S*.Tm that detach, invade, or remain on the IEC surface are quantified for n=3 replicate time-lapse movies for each condition. Data in B-C derive from donor 1 enteroid cultures and donor 4 colonoid culture experimentation. The results revealed a dramatically reduced permissiveness to *S*.Tm invasion for ENR-grown enteroid-and colonoid-derived monolayers. **(D)** Similar-sized HeLa cell monolayers, IEC monolayers grown in OGM, or in ENR, were infected with 10^7^ CFUs of *S*.Tm for 40min. Shown are the CFU counts of *S*.Tm *wt* intracellular bacteria after gentamicin treatment. Statistical analysis; Wilcoxon rank-sum test, unpaired. *p-value = 0.048. **(E-F)** Quantification of *S*.Tm invasion foci by a 96-well-format semi-automated imaging assay. Enteroid-derived 2D monolayers were grown in OGM or ENR within microwells, followed by infection with an *S*.Tm *wt*/p*ssaG*-GFP reporter strain at MOI 40 for 20min, followed by 3h further incubation for reporter maturation. **(E)** Representative fluorescence micrographs of *S*.Tm *wt*/p*ssaG*-GFP-infected monolayers stained with DAPI (nuclei) and phalloidin (F-actin). Scale bars, 50µm. **(F)** Quantification of the number of *S*.Tm invasion foci (*ssaG*-GFP+) per field of view for infected OGM and ENR-grown monolayers. Also in this assay, the ENR monolayers were less permissive to *S*.Tm invasion that their OGM counterparts. Data pooled from 4 independent experiments presented with one infected well as one data point (n=10 and n=11, respectively). Statistical analysis; Wilcoxon rank-sum test, unpaired. ***p-value = 0.00012.

The structural host cell alterations coupled to *S*.Tm invasion may vary from expansive lamellipodia-and filopodia-containing ruffles (seen in flat-growing tumor cell lines) to more discreet and transient entry structures (seen e.g. in the mouse gut) (10). Therefore, we wanted to validate the present findings by a method not relying on time-lapse imaging. For this purpose, gentamicin-protection assays were performed for OGM and ENR monolayers, using the well-characterized HeLa cell line as an additional reference. The results supported the conclusions from live-cell imaging. After exposure to the same sized *S*.Tm inoculum (10^7^ CFUs, infection for 40min) enteroid-derived ENR monolayers contained ∼8-fold fewer intracellular bacteria than the corresponding OGM monolayers, and approximately 100-fold fewer than an equally-sized cell layer of the HeLa reference culture (**Fig 2D**). As a third analysis type, we developed a 96-well format infection assay, where IEC monolayers were grown in OGM or ENR within microwells, followed by infection with an *S*.Tm *wt*/p*ssaG*-GFP reporter strain (turns GFP+ upon arrival in the *Salmonella*-containing vacuole (45, 46)). This permitted scoring of the number of successful invasion events by a semi-automated image analysis pipeline. Again, IEC monolayers cultured in OGM medium contained more GFP-focus-positive IECs than the corresponding ENR monolayers (**Fig 2E-F**). Taken together, these data show that differentiation of human IECs from an immature stem/progenitor cell state (OGM) towards an enterocyte or colonocyte state (ENR) drastically reduces the permissiveness to *S*.Tm invasion.

### Characteristics of the *Salmonella* Invasion Process in Immature versus Maturing Human Intestinal Epithelial Cell Layers

*S*.Tm invasion of an epithelial cell is fueled by delivery of TTSS-1 effectors that manipulate actin-regulatory networks, including Rho-and Arf-GTPases and formins. This drives actin polymerization and bundling and plasma membrane deformation (47). We examined if differential expression of actin regulators might explain the differing permissiveness to *S*.Tm invasion between immature and maturing IEC monolayers. Global proteomic analysis was conducted on enteroid-derived monolayers grown in OGM, ENR, or a commercial equivalent of ENR (StemCell Organoid Differentiation Medium, “cENR”) medium. This resulted in the identification of 7328 proteins, and quantification of 5978 proteins across samples based on the detection of at least two unique and razor peptides. A clear separation of the proteomes was observed between 3D enteroids in OGM (used as additional control), OGM monolayers and ENR/cENR monolayers (clustered closely together) (Principal component analysis and Venn diagrams in **Fig S3A**). Mining of the proteomes from ENR/cENR compared to OGM monolayers revealed lower levels of GTPases (CDC42, RHOA, ARF1; (11, 12, 14, 48)) and formins (FHOD1; (15)) previously implicated in *S*.Tm invasion (**Fig 3A-C**).

**Figure 3.**
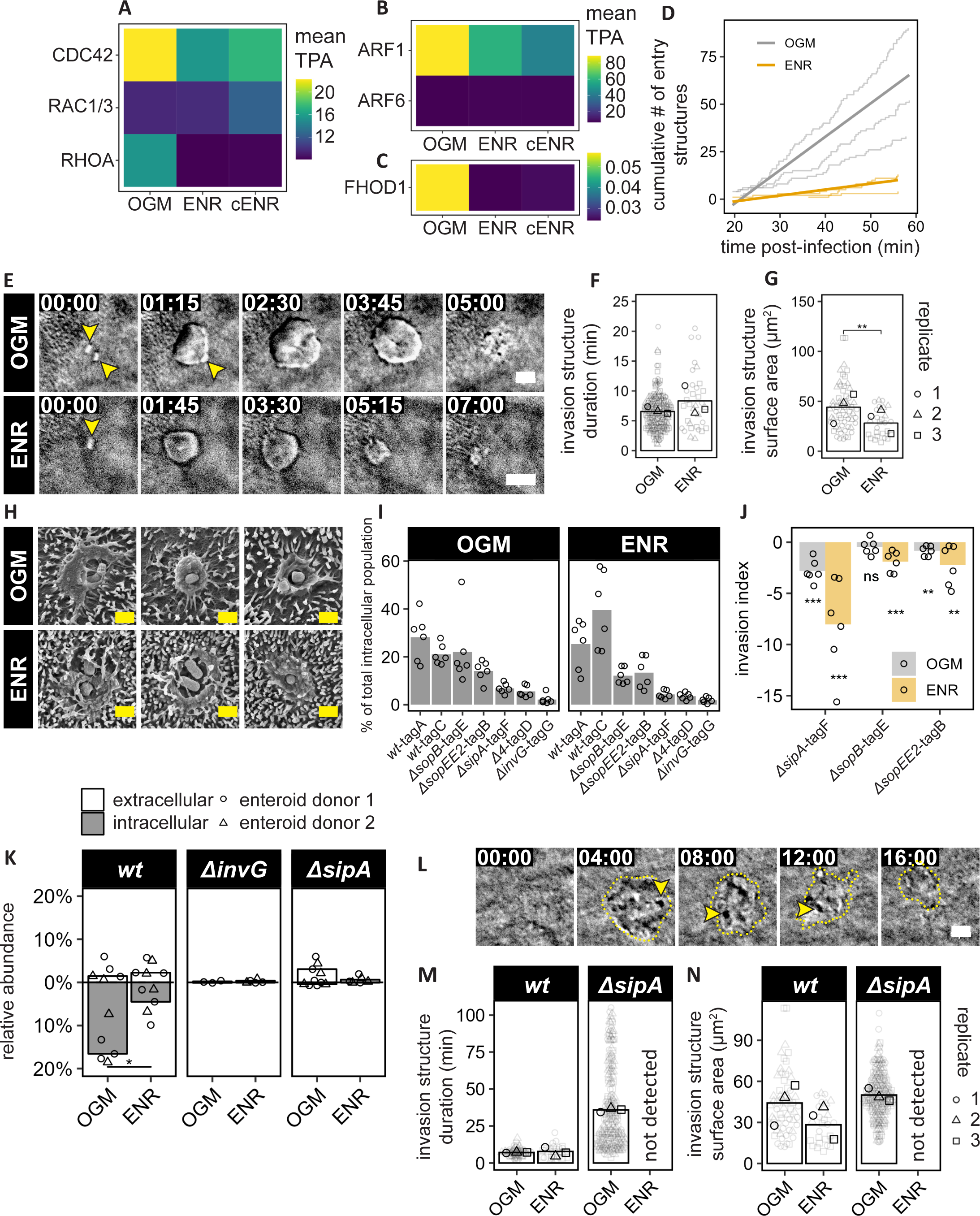
The *Salmonella* invasion step in human intestinal epithelia of different maturity. **(A-C)** Quantitative proteomic profiling of actin regulatory proteins in enteroid-derived IEC monolayers grown in OGM, ENR, or commercial ENR (cENR). Heatmaps show absolute protein quantitation using the total protein approach (TPA; femtomole per microgram total protein) for each indicated protein. Mean values from n=3 biological replicates per condition shown for select **(A)** Rho GTPases, **(B)** Arf GTPases, and **(C)** a formin detected in the proteomes. The results showed a trend towards lower expression of these actin regulatory proteins, previously implicated in *S*.Tm host cell invasion, upon human IEC maturation. **(D-G)** Time-lapse imaging of enteroid-derived IEC monolayers grown in OGM or ENR within AICs and infected with *S*.Tm *wt* at MOI 3 for 1h. **(D)** Quantification of the cumulative number of entry structures detected over time. **(E)** Representative time-lapse DIC image series of *S*.Tm-elicited entry structures (yellow arrowheads) at the apical surface of OGM-(top row) and ENR-grown (bottom row) monolayers. Time in min:sec; scale bars, 5µm. Quantification of entry structure **(F)** duration and **(G)** maximal area for time-lapse movies as in E. Each symbol represents one entry structure pooled from 3 replicate time-lapse movies for each condition. Statistical analysis; Kruskal-Wallis test followed by Dunn’s post-hoc with Holm-Bonferroni correction (as an additional group was also part of the dataset; see Fig S3E). **p-value = 0.00819. **(H)** Representative scanning electron microscopy images of *S*.Tm-elicited apical entry structures in OGM and ENR monolayers. Scale bars, 1µm. The presence of some elongated “doublet bacteria” among the invading *S*.Tm has been explained in earlier work (9). Taken together, the results from D-H showed a drastic reduction in the frequency of *S*.Tm-elicited entry structures between OGM and ENR monolayers, and a modest decrease in average entry structure size. **(I-J)** Apical-out 3D enteroids grown in OGM or ENR were infected with a mixed barcoded consortium comprising the indicated strains at MOI 40 for 20min. Shown is the relative abundance of the indicated *S*.Tm strains in the intracellular population, normalized on the corresponding input, as quantified by **(I)** qPCR, and with **(J)** the respective invasion indexes (see Methods for calculation). Bars correspond to means of six replicate infections pooled from two different occasions. Each replicate is indicated by a circle symbol. Statistical analysis; paired t-test between input - output pairs. ns p-value = 0.75, **p<0.01, ***p<0.001. **(K)** Quantification of intra-and extracellular bacteria in enteroid-derived monolayers grown in OGM or ENR on PET transwell supports, infected by *S*.Tm *wt*, *`Δ`invG*, or *`Δ`sipA* strains carrying the *rpsM*-mCherry constitutive reporter at MOI 3 for 1h, and subsequently fixed and stained for anti-*Salmonella* LPS without permeabilization. The y-axis shows the abundance of the intra-(negative) or extracellular (positive) bacteria normalized across strains and conditions (see Methods for details). Each symbol represents one replicate for donor 1 (n=3) or 2 (n=2), bars indicate the median. These findings substantiated a lower frequency of *S*.Tm *wt* invasion events in ENR than in OGM monolayers, and a profound impacts of SipA for successful IEC invasion in either condition. Statistical analysis; Wilcoxon rank-sum test, unpaired. *p-value = 0.016. **(L-N)** Time-lapse imaging of enteroid-derived monolayers grown in OGM or ENR within AICs and infected with *S*.Tm *wt* or *`Δ`sipA* at MOI 3 for 1h. **(L)** Representative time-lapse DIC image series blow-up of a *S*.Tm *`Δ`sipA*-elicited entry structures (yellow dashed outline) at the apical surface of a OGM monolayer. Time in min:sec; scale bars, 5µm. Quantification of entry structure **(M)** duration and **(N)** maximal area for time-lapse movies as in E. Each symbol represents one entry structure pooled from 3 replicate time-lapse movies for each condition. The *S*.Tm *wt* data are the same as presented in panels F-G. The data in L-N provided further evidence for the key importance of *`Δ`sipA* for *S.*Tm invasion of ENR monolayers. Moreover, the *S*.Tm *`Δ`sipA* strain elicited prolonged “frustrated ruffling” in OGM monolayers which failed to convert into successful invasion events.

The proteomics results hinted that the propensity for *S*.Tm-effector-induced membrane ruffling may decrease during human IEC maturation. Indeed, vastly fewer entry structures were visible by live-cell imaging in the ENR condition (**Fig 3D**, see also **Fig 2A-C** above). Morphological analysis of the detectable entry structures showed broadly similar characteristics between conditions. In live-cell DIC microscopy, entry structures had a donut-like or interdigitated appearance and were transient in nature (**Fig 3E-F**). We quantified the mean entry structure duration to ∼7-8min in both OGM and ENR monolayers (**Fig 3F**), whereas the maximal entry structure area was significantly smaller in the ENR monolayers (28.21 ± 13.69µm^2^ vs 44.17 ± 23.08µm^2^ for OGM monolayers; **Fig 3G**). SEM of *S*.Tm invasion events captured mid-way supported these conclusions and illustrated that IEC surface perturbations were confined to the area immediately close to the bacterium (**Fig 3H**). Parallel analysis of the additional ENRRT (M-cell-like) control sample again revealed more frequent and larger entry structures than for ENR monolayers (**Fig S3C-F**).

To assess the quantitative impact of *S*.Tm TTSS-1 effectors, we next adapted barcoded consortium infections to infect human enteroids with multiple mutant and wild-type *S*.Tm strains (20). To enable bulk infections and avoid bottleneck effects due to low total bacterial population sizes, we generated apical-out 3D enteroids in liquid culture and maintained these in either OGM over 3 days or ENR medium over 6 days (see Methods for details (32)). The enteroids were subsequently infected from the apical side with a barcoded consortium containing a 1:1:1:1:1:1:1 mix of the seven *S*.Tm strains: *wt*:tagA, *wt*:tagC (wild-type controls), *ΔsopB*:tagE (lacks the SopB effector), *ΔsopEE2*:tagB (lacks SopE and SopE2), *ΔsipA*:tagF (lacks SipA), *Δ4*:tagD (lacks SopB, SopE, SopE2, and SipA), and *ΔinvG*:tagG (lacks a key structural component of the TTSS-1 (20)). qPCR-based detection of the unique chromosomal tags allowed quantification of each strain’s abundance in the inoculum and in the bacterial population successfully invading the enteroids (49). An analogous setup recently showed that *S*.Tm invasion of flat-growing epithelial cell lines (HeLa) heavily depends on SopB/E/E2 with a neglectable impact of SipA, while in the mature intestinal epithelium of the mouse gut, SipA is of key importance (10). Here, we in human enteroids found that strains deleted for SopB, or SopE and E2, were partially attenuated, but that deletion of SipA again resulted in the strongest invasion defect (**Fig 3I-J**). This was evident in apical out OGM enteroids, but even more pronounced upon enteroid maturation in the ENR medium (**Fig 3I-J**). The strong dependence on SipA for human IEC invasion was further substantiated in IEC monolayers infected with constitutively fluorescent *S*.Tm strains followed by anti-LPS staining to resolve extracellular and intracellular bacteria (**Fig 3K**). Finally, we again employed live-cell imaging in AICs to compare the entry structure characteristics elicited by *S*.Tm *wt* versus *S*.Tm *ΔsipA*. This led to two notable observations. First; no visible entry structures at all were detected by DIC imaging of maturing ENR monolayers infected with *S*.Tm *ΔsipA*, which was in sharp contrast to *S*.Tm *wt* infections (**Fig 3L-N**). Second; in the immature OGM monolayers, *S*.Tm *ΔsipA* bacteria did elicit entry structures, but these had a dramatically prolonged duration (35.87 ± 26.88min, compared to 7.06 ± 2.83min for *S*.Tm *wt*) where the eliciting bacterium floated along the surface, but failed to complete invasion into the IEC (**Fig 3L-M**). From these and earlier data, we conclude that SipA is of particular significance for *S*.Tm invasion of mature non-transformed polarized murine and human IECs (Fig 3) (10), and that this effector promotes conversion of TTSS-1-elicited entry structures into productive invasion vessels (Fig 3) (10, 50).

In summary, maturation of human IECs from an immature progenitor (OGM) state towards a mature enterocyte (ENR) state is coupled to i) modestly lowered expression of actin-regulators implicated in *S*.Tm-induced ruffling, ii) a ∼36% reduction in entry structure size, and iii) an increased dependence on the TTSS-1 effector SipA for successful *S.*Tm invasion. However, these differences connected to the actual invasion process itself appeared insufficient to explain the drastically lower total number of *S*.Tm successfully attacking the ENR monolayers, compared to the OGM counterparts (**Fig 2**).

### Human Intestinal Epithelial Cell Maturation towards an Enterocyte/Colonocyte State Fortifies the Apical Surface against *Salmonella* Attack on the Membrane

The host cell invasion step is preceded by *S*.Tm reaching the IEC surface and inserting the TTSS-1 translocon into the membrane for effector delivery. By live-cell imaging, we had noted a considerable lag time between the binding of the bacteria atop the IEC surface and initiation of invasion (**Fig 2**; (26)). This “surface lingering” was particularly apparent in ENR monolayers and may suggest that human IEC maturation fortifies an apical microbarrier that *S*.Tm has to overcome. Proteome analysis indeed revealed higher levels of e.g. several intermicrovillar adhesion complex proteins (CDHR2, CDHR5, MYO7B, USH1C), the surface-attached mucins MUC1 and MUC13, the galactosyl transferase B3GALT5, and the fucosyl transferases FUT2 and FUT3, in ENR/cENR than in OGM monolayers (**Fig 4A-C**). The major soluble mucin MUC2 was as expected at or below the detection limit (**Fig 4B**). Surface-attached mucin proteins like MUC1 and MUC13 comprise elongated and heavily O-glycosylated components of the apical IEC glycocalyx (24). They have been implicated in barrier functions while at the same time serving as potential ligands for bacterial adhesins (7, 51–53). Lectin stainings of fixed samples further demonstrated shifts in the apical glycan composition between OGM and ENR conditions (**Fig 4D-E**). Specifically, IEC monolayer maturation resulted in a lowered mean signal for apically exposed GlcNAc (detected by Wheat-germ agglutinin; WGA), but elevated signals for both terminal fucose and sialic acid glycans (detected by *Ulex europaeus* agglutinin-I; UEA-I, and *Sambucus nigra* lectin; SNA-I, respectively) (**Fig 4D-E**) (54). Of note, all three lectins also highlighted substantial cell-to-cell heterogeneity both for OGM and ENR monolayers (**Fig 4D-E**).

**Figure 4.**
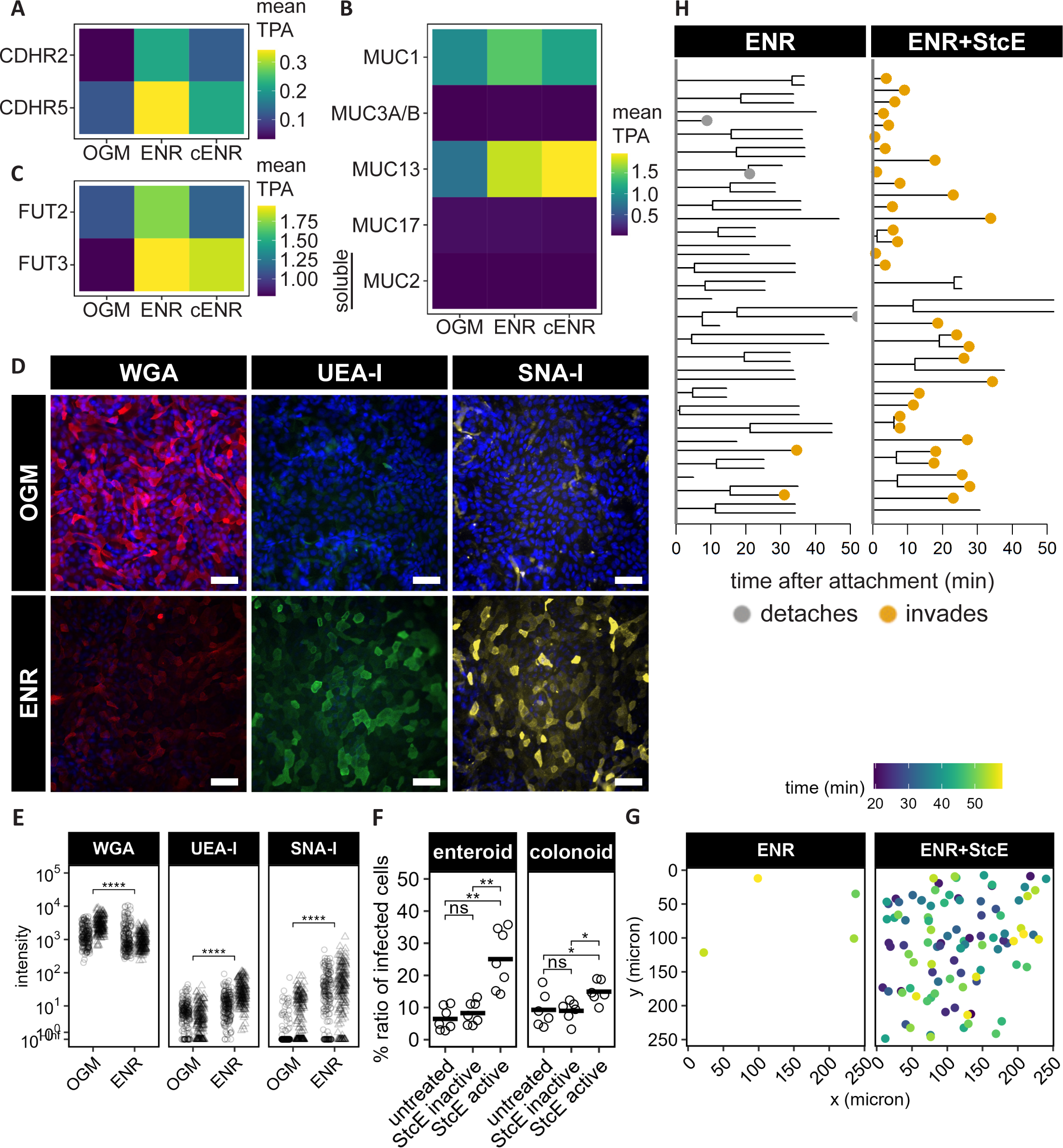
Human intestinal epithelial cell maturation towards an enterocyte/colonocyte phenotype fortifies the apical surface against *Salmonella* invasion. **(A-C)** Quantitative proteomic profiling of proteins relevant to the apical IEC surface in enteroid-derived monolayers grown in OGM, ENR, or commercial ENR (cENR). Heatmaps show absolute protein quantities using the total protein approach (TPA; femtomole per microgram total protein) for each indicated protein. Mean values from n=3 biological replicates per condition are shown for select **(A)** microvillus-crosslinking cadherins, **(B)** mucins, and **(C)** fucosyl transferases detected in the proteomes. The results revealed robust upregulation of e.g. CDHR5, FUT3, and MUC13 upon human IEC maturation. **(D-E)** Quantification of the apical IEC surface glycocalyx composition by a panel of lectins. **(D)** Representative micrographs of 2D enteroid-derived monolayers grown in OGM (top row) or ENR (bottom row), fixed and stained with DAPI (nuclei) and the lectins WGA (detects GlcNAc), UEA-I (detects fucose), or SNA-I (detects sialic acid). **(E)** The mean fluorescence intensity per cell was quantified in 200 randomly selected IECs per field of view and the background from unstained samples subtracted. Data pooled from two independent experiments, showing results from two independent enteroid cultures (donor 1 and donor 2) side-by-side. These data demonstrated decreased average WGA signal and increased average UEA-I and SNA-I signals, upon IEC maturation. A substantial cell-to-cell heterogeneity was also observed for all conditions. Statistical analysis; Wilcoxon rank-sum test, unpaired. **** p<0.0001. **(F-H)** Quantification of the impact of StcE pretreatment on enteroid-derived IEC monolayer permissiveness to *S*.Tm invasion. **(F)** Enteroid-and colonoid-derived IEC monolayers were grown in ENR within microwells, left untreated, treated for 2h with enzymatically inactive StcE, or active StcE, and subsequently infected with the *S*.Tm *wt*/p*ssaG*-GFP reporter strain. The graph depicts the percentage of infected (*ssaG*-GFP+) IECs following infection at MOI 40 for 20min, followed by 3h further incubation for reporter maturation. StcE pretreatment led to significantly increased numbers of *S*.Tm invasion foci in both enteroid-and colonoid-derived ENR monolayers. Statistical analysis; Kruskal-Wallis followed by Dunn’s post hoc test with Benjamini-Hochberg correction. *p<0.05, **p<0.01. **(G-H)** Time-lapse imaging of enteroid-derived 2D monolayers grown in ENR within AICs, left untreated or treated with StcE, and subsequently infected with *S*.Tm *wt* at MOI 3 for 1h. In **(G)** spatiotemporal maps of detected *S.*Tm invasion events over the field of view is shown. In **(H)** the behavior of individual surface-bound *S*.Tm is visualized as dendrograms for each condition. The dendrograms indicate the timing of bacterial division (track split), detachment (grey dot), invasion (yellow dot), and remaining bacteria at the end of the movie or loss of focus (bare branch). This analysis corroborated that StcE pretreatment converts ENR-grown monolayers back to a *S*.Tm-invasion-permissive phenotype.

To examine if these apical surface alterations could explain ENR monolayer fortification against *S*.Tm, we made use of the bacterial enzyme StcE, previously shown to cleave the protein backbone of surface-attached mucins in a glycosylation-dependent manner (55, 56). We reasoned that if the fortification of the maturing (ENR) monolayers stemmed from the build-up of the apical mucin-linked glycocalyx, then StcE pretreatment should revert these monolayers back to the *S*.Tm-invasion-permissive phenotype of the OGM counterparts. We first employed the 96-well *S*.Tm invasion assay to test this hypothesis. Strikingly, pretreatment of enteroid-derived ENR monolayers with enzymatically active StcE, but not a catalytically dead StcE variant, led to a >3-fold increase in the number of successful *S*.Tm invasion (*ssaG*-GFP+) events (**Fig 4F, Fig S4A**). This also held true for colonoid-derived ENR monolayers (**Fig 4F**). As death of many infected IECs already become evident within the 3h 20min time frame of this assay (**Fig S4B**), we finally turned to time-lapse imaging in AICs to visualize *S*.Tm invasion events in real-time during the first hour of the infection (preceding signs of IEC death). This again revealed only sparse *S*.Tm invasion events in untreated ENR monolayers. Remarkably, however, StcE pretreatment fueled a >10-fold increase in the number of visible *S*.Tm invasion events, spaced out evenly over the monolayer (**Fig 4G-H**). Altogether, these results demonstrate that human IEC differentiation from an immature progenitor state towards a maturing enterocyte/colonocyte state fortifies the apical surface against *S*.Tm attack.

## Discussion

Our mechanistic understanding of how pathogens invade IECs has been gained largely from cell line infections. Recent work has, however, indicated that although *S*.Tm invades both flat-growing epithelial cell lines and the intact murine gut absorptive epithelium in a TTSS-1-dependent manner, there are striking differences. *S*.Tm invasion of murine gut IECs appears significantly more challenging for the bacterium (a higher effective MOI and longer time is required), involves smaller entry structures, and relies on TTSS-1 effector activities in different proportions than seen in cell lines (10). Comparative microscopy has further illustrated that *S*.Tm swiftly invades HeLa and Caco-2 cells within minutes of reaching the cell surface, but that in human enteroid-derived epithelia the same process involves a step where the bacteria linger atop the surface for prolonged periods before managing to invade (26). The present study begins to explain these discrepancies. First, human enteroid-and colonoid-derived epithelial cell layers remain permissive to *S*.Tm invasion when maintained in an immature state, whereas maturation towards an enterocyte or colonocyte phenotype dramatically enhances IEC layer resistance to the attack. Second, human IEC maturation involves shifts in expression of actin regulators usurped by the pathogen. Third, the maturation process includes build-up of the apical surface mucin-linked glycocalyx, which limits bacterial access. These findings demonstrate how the maturation state of human host epithelial cells fundamentally impacts their permissiveness to invasion by a model enteropathogen, thus reconciling apparent previous contradictions in the literature.

The combined activities of four main effectors, namely SopB, SopE (present in *S*.Tm SL1344, but not in e.g. *S*.Tm 14028), SopE2, and SipA drive TTSS-1-dependent *S*.Tm invasion of epithelial cells both in cell line culture, enteroids, and *in vivo* mouse models (10, 20, 27, 57–59) (and this work). Among these effectors, SopB/E/E2 directly or indirectly activate Rho GTPases (11–13, 60), Arf GTPases (14), and formins (15) to fuel actin polymerization and expansive membrane ruffle formation. SopB/E/E2 activity suffices to accomplish wild-type levels of *S*.Tm invasion into non-polarized cell lines, whereas deletion of SipA has a negligible effect (10, 20). This stands in stark contrast to findings from both adult and neonate mice, where SipA has been assigned an influential role (10, 58). The present study generalizes these latter findings, as we detect a marked invasion defect of SipA mutant *S*.Tm in non-transformed human IEC layers, and even more so following their maturation. Hence, we propose that as mammalian IECs mature towards an enterocyte or colonocyte phenotype (and away from the immature phenotype most evident for tumor cell lines), their propensity to permit SopB/E/E2-driven ruffling gradually diminishes. Instead the importance of SipA-mediated local actin stabilization (17–19, 61) for successful IEC invasion increases (10) (and this work). It is noteworthy that our global proteomic analysis revealed lower levels of several established SopB, E and E2 (direct or indirect) targets, including Cdc42, RhoA, Arf1, and FHOD1 (11, 14, 15, 48), following human IEC maturation.

Time-lapse DIC and SEM imaging illustrated the presence of apical *S*.Tm entry structures in both immature OGM and maturing ENR IEC monolayers, but with a markedly lower frequency, and a ∼30% reduced surface area, in the latter. Since also the maturing IEC layers remain heterogeneously differentiated, it is possible that the entry structures still observed under this condition preferentially arise in relatively immature cells. Nevertheless, entry structures were as a rule small and transient with a mean duration of ∼7-8 minutes, hence differing from the expansive ruffles elicited by *S*.Tm in e.g. HeLa cells (10, 22, 62), but with some resemblance to the corresponding structures seen in polarized MDCK cells (10, 50, 63). Further in line with pioneering work in polarized MDCK (50), an *S*.Tm strain lacking the SipA effector elicited strikingly long-lived apical membrane structures (mean duration of >30 minutes) in immature IEC layers, but which failed to convert into productive vessels for bacterial invasion. In maturing IEC layers, *S*.Tm invasion was essentially abrogated in the absence of SipA. Taken together, we conclude that apical membrane ruffling downstream of *S*.Tm TTSS-1 effectors is constrained by enterocyte/colonocyte maturation, thereby providing an obstacle to bacterial invasion. Still, this effect on its own is insufficient to account for the >10-fold drop in *S*.Tm invasion frequency noted between immature and maturing absorptive IEC layers.

Proteomic and morphological assessment further revealed changes in the apical surface ultrastructure and composition during epithelium maturation. This includes microvillus densification, elevated expression of microvillus-crosslinking cadherins and the cell surface mucins MUC1 and MUC13. It should be noted that mucins are typically poorly cleaved and detected in global proteomics pipelines. Hence, the current quantifications of MUC1, MUC13, and possibly other related mucins, likely represent underestimates. Cell surface mucins constitute the backbone for the apical IEC glycocalyx (24). In the case of MUC1 and MUC13, these glycoproteins have been implicated as both ligand structures for bacterial adhesins, and barriers to diverse microbes (7, 51–53). The elevated expression of fucosyl transferases, and increase in lectin-stainable fucose and sialic acid-containing glycans also substantiate that the switch between our OGM (immature) and ENR (maturing) conditions causes glycan maturation (64). Most importantly, we found that enzymatic “shaving” of the apical surface by the selective mucinase StcE (55, 56) reverts maturing human IEC layers back to the *S*.Tm-invasion-permissive phenotype of the immature counterparts. This provides a causal link between the buildup of a complex mucin-linked glycocalyx during human enterocyte/colonocyte maturation and a reduced permissiveness to *S*.Tm invasion. These findings gain independent support from the observations that MUC13 knockout mice are hypersusceptible to *S*.Tm gut infection, and that MUC13 can in principle function as both a physical barrier and a bacterial binding decoy (51). At the same time, our data do not refute that *S*.Tm binding of MUC1 as a ligand for the giant SiiE adhesin can under some specific experimental conditions promote invasion (7). The biochemical structure -function relationships that explain the apical IEC glycocalyx barrier properties and its intersection with bacterial virulence factors remains an exciting area for future research.

Despite its obvious advantages over traditionally used cell lines, our experimental model system has its limitations. These human 2D IEC layers are in the present versions devoid of a soluble mucus layer, lack luminal flow and an accurate representation of 3D crypt - villus geometry. Moreover, the culture protocols applied only partially direct differentiation towards a true enterocyte, colonocyte or M-cell end state. This supports that chemical, physical, and/or geometrical cues originating from the native tissue context or the microbiota, are needed to fuel terminal IEC differentiation (65, 66). For example, IEC expression of the long surface mucin MUC17 has been shown to require interleukin-22 and potentially other signals from the lamina propria (67), and is also promoted by villus geometry (68). This mucin remains close to undetectable on the protein level in our enteroid monolayer cultures, suggesting that the here identified barrier effect to *S*.Tm infection may be even more potent *in vivo*. Nevertheless, our study provides a foundation for the systematic reconstruction of the human intestinal epithelial infection niche under conditions compatible with high resolution live-cell imaging. We anticipate this and similar approaches to have broad implications for the merger of biochemical mechanism and host cell and tissue physiology in the new era of infection research.

## Methods

### Ethics statement

Human enteroids were either established in previous studies (26–28), or enteroids and colonoids established *de novo* here. Enteroids were established from jejunal tissue resected during bariatric surgery. Colonoids were established from morphologically normal non-tumor tissue resected as margin during elective colon cancer surgery. All tissue samples were acquired following informed written consent. Samples were pseudonymized prior to further processing, so that the patients’ identities and personal information remained unknown to researchers working with the samples. For this study, enteroid cultures Hu18-8jej, Hu18-9jej, and Hu22-2jej, and colonoid cultures Hu21-9col and Hu22-4col were used. The procedures were approved by the local governing body (Etikprövningsmyndigheten, Sweden) under license numbers 2010-157, 2010-157-1, 2020-05754, and 2023-01524-01.

### Human enteroid and colonoid establishment

*De novo* establishment of enteroids and colonoids was performed essentially as described for enteroids before (27). Briefly, resected intestinal tissue samples were cut into millimeter-sized pieces, washed in ice-cold phosphate-buffered saline (PBS) (#12559069, Thermo Fisher/Gibco, Waltham, MA, USA) and epithelial crypts subsequently dissociated using gentle cell dissociation reagent (#100-0485, STEMCELL Technologies, Vancouver, BC, Canada) and nutation at 4°C for 30min in case of jejunal samples, and 45min in case of colon samples, followed by trituration. The resulting epithelial fragments were filtered through a 70µm cell strainer (#15370801, Corning, Corning, NY, USA) and crypt-enriched fractions resuspended in 50µl Matrigel (#356230, Corning, Corning, NY, USA) domes in 24-well plates (#83.3922, Sarstedt, Nümbrecht, Germany). The embedded crypts were cultured in OGM (IntestiCult Human organoid growth medium; #06010, STEMCELL Technologies) with 100U/ml penicillin-streptomycin (#15140122, Thermo Fisher/Gibco) at 37°C and 5% CO_2_. The OGM medium was refreshed every 2 to 3 days. Newly established enteroids and colonoids were cryopreserved at passage 1-4 by gently dissolving Matrigel (Corning) domes in ice-cold Dulbecco’s modified Eagle’s medium DMEM/F12 (#11580376, Thermo Fisher/Gibco) with 0.25% bovine serum albumin (BSA; #11500496, Thermo Fisher/Gibco) and resuspending the enteroids or colonoids in DMEM/F12/10% heat-inactivated fetal bovine serum (FBS; #11573397, Thermo Fisher/Gibco)/10% DMSO (#D2438, Sigma-Aldrich, Stockholm, Sweden), followed by freezing in a Mr. Frosty^TM^ freezing container at -80°C overnight and transfer to liquid nitrogen gas phase for long-term cryopreservation.

### Enteroid and colonoid culture

For maintenance of recently established and thawed enteroids and colonoids, the cultures were passaged weekly at a ratio of ∼1:8 for enteroids, or ∼1:6 for colonoids. The Matrigel domes were manually broken up by pipetting with gentle cell dissociation reagent and then washed once with DMEM/F12/0.25% BSA. The resulting resuspended enteroids/colonoids were disrupted by triturating 15-20 times with a 200µl pipette tip. Following disruption, fragments were again resuspended in 50µl domes of Matrigel:OGM at a ratio of 3:1, divided over 3 domes per well in 24-well plates, and cultured at 37°C and 5% CO_2_. The OGM medium was refreshed every 2 to 4 days.

### Enteroid-and colonoid-derived IEC monolayer culture

Human IEC monolayers were cultured on either 24-well transparent PET tissue culture transwell inserts with 0.4µm pores (#83.3932.041, Sarstedt, Nümbrecht, Germany), 13-mm-diameter alumina Whatman Anodisc membranes (#WHA68097023), or in 96-well plates (tissue culture treated, sterilized, #89626, ibidi). The PET transwell inserts and 96-well plates were coated with 40x diluted Matrigel in PBS for 1h at 37°C prior to use. After this, the coating solution was completely removed and the cell suspension immediately added to the transwell inserts or wells. The Anodisc alumina membranes were coated by a sequential protocol, as further detailed in (26). First, the alumina membranes were soaked in 30% H_2_O_2_ for 1h at room temperature, followed by washing in sterile distilled water (dH_2_O) and incubation in 0.1mg/ml poly-L-lysine (#P6282, Sigma-Aldrich) in dH_2_O for 5min. The membranes were then air dried in a laminar-flow cabinet for >2h. Finally, the membranes were soaked in 40x diluted Matrigel in dH_2_O for 1h at 37°C and air dried again. Coated membranes were mounted within custom-manufactured Apical Imaging Chambers (AICs (26); design files available at https://doi.org/10.17044/scilifelab.16402539). For monolayer culturing, circa one well of 3D enteroids/colonoids kept in OGM per membrane was dissociated into single cells at day 7 to 9 after passaging. The enteroids/colonoids were first taken up from the Matrigel domes in gentle cell dissociation reagent and then washed in DMEM/F12/1.5%BSA and dissociated into single cells using TrypLE Express (#10043382, Thermo Fisher/Gibco) for 5-10min at 37°C. Cells were then spun down at 300g, 4°C, for 5min and resuspended in OGM/10µM Y-27632 (ROCK inhibitor; #72304, STEMCELL Technologies). Cells were enumerated in a Bürker chamber, and 2.5x10^5^ cells were seeded into the apical compartment of PET transwell inserts in 150µl (600µl medium in the basolateral compartment, 24-well plate wells), 2.5x10^5^ cells per well of a 96-well plate in 200µl, or 3.0x10^5^ cells into the apical compartment of AICs in 100µl (500µl medium in the basolateral compartment, kept in 12-well plate wells). The monolayers grew confluent within ∼3 days (range 2-4). For a stem cell/progenitor IEC phenotype, monolayers were maintained in OGM until analysis on days 3-7. For monolayer maturation towards and enterocyte/colonocyte phenotype, the medium was replaced with ENR (EGF-Noggin-R-spondin1-containing, but lacking Wnt3a). This medium is based on DMEM/F12 supplemented with 5% R-Spondin1-conditioned medium (made from Cultrex 293T R-spondin1-expressing cells; R&D Systems, MN, USA), 10% Noggin-conditioned medium (made with HEK293-mNoggin-Fc cells; kindly provided by Hans Clevers, Utrecht University), 50ng/ml mouse recombinant EGF (#SRP3196, Sigma-Aldrich), 1xB27 supplement (#11530536, Gibco), 1.25mM N-acetyl cysteine (#A7250, Sigma-Aldrich), and 100U/ml penicillin-streptomycin. For IEC differentiation towards an M-cell-like phenotype, ENRRT medium was used, generated by supplementing the ENR medium with 200ng/ml Recombinant Mouse TRANCE (RANKL) (#577102, BioLegend) and 50ng/ml Recombinant Human TNF (#PRSI40-115, ProSci). In some experiments we also used an additional commercially available ENR medium (here denoted cENR; IntestiCult Human organoid differentiation medium; #100-0214, STEMCELL Technologies) with 100U/ml penicillin-streptomycin. Monolayer differentiation was allowed for ∼4 additional days (range 4-5 days) after reaching confluence. For infection experiments, the medium was in all cases replaced with DMEM/F12 without antibiotics on the day of experimentation.

### Transcriptome analysis

Enteroids from two donors were grown as 3D cultures in OGM or as monolayers in AICs in OGM, ENR, and ENRRT. For transcriptomic analysis using RNA-sequencing, we isolated total RNA from these samples using Trizol LS (Invitrogen) according to the manufacturer’s protocol. Briefly, two wells of 3D organoids or one disassembled Anodisc membrane were washed and lysed with 500µl Trizol, and RNA was extracted with 100 µl chloroform by manual shaking. The Trizol/chloroform mixture was then spun down and RNA was precipitated from the aqueous phase using isopropanol. The RNA was spun down into a pellet, washed with 80% ethanol, air dried, and resuspended in 20µl RNase-free water. The RNA concentration was measured with a Nanodrop (Thermo Fisher), and ≥1µg per sample was submitted to the National Genomics Infrastructure Sequencing Facility of SciLifeLab, Uppsala. There, library preparation was performed using the TruSeq stranded mRNA (Illumina) protocol with poly-A selection and sequenced using one lane of a S4-300 chip on a NovaSeq 6000 with a paired-end (2x150 bp) setup. Alignment was performed by the facility. TPM values were calculated based on final gene counts in R to generate a Principal Component Analysis (Fig S1) and a curated gene heatmap (Fig 1B).

### Proteome analysis

3D enteroids and enteroid-derived IEC monolayers were grown as described above in either OGM, ENR, or cENR (without DAPT). Three biological replicates of each sample type were used for global proteomics analysis (69). Samples were collected, lysed (50 mM DTT, 2 %(w/v) SDS, 100mM Tris/HCl pH 7.8) and MED-FASP was performed using Trypsin and Lys-C enzymes. C18 stage tips were used to desalt the peptide mixture and samples were stored at −20°C until analysis. Protein and peptide content were determined by the tryptophan fluorescence assay (70). The global proteomics analysis was performed on a Q Exactive HF mass spectrometer (MS) (Thermo Fisher Scientific) coupled to a nano–liquid chromatography (nLC). An EASY-spray C18-column (50cm long, 75μm inner diameter) was used to separate peptides on an acetonitrile/water gradient with 0.1% formic acid over 150min. MS was set to data dependent acquisition with a Top-N method (full MS followed by ddMS2 scans). Proteins were identified using the MaxQuant software (version 2.1.0.0) with the human proteome reference from UniProtKB (April, 2022). The total protein approach was used as protein quantification method (71).

### *S.*Tm strains, plasmids, and inoculum preparation

All *S.*Tm infections were performed with strains of the *Salmonella enterica* serovar Typhimurium SL1344 background (SB300, streptomycin resistant; (72). Besides the wild-type (*wt*), the study employed previously characterized *ΔsipA* and *ΔinvG* strains for microscopy (10, 73). Barcoded *S*.Tm *wt* and mutant strains, carrying a unique genetic 40-nt tag within the *S*.Tm malXY locus have also been described and validated before (20, 49). For fluorescence microscopy applications, the indicated *S*.Tm strains carried either a pFPV-mCherry reporter plasmid (*rpsM*-mCherry; Addgene plasmid number 20956; (74)) directing constitutive mCherry expression, or the p*ssaG*-GFP reporter plasmid (pM975; (45, 46)) directing GFP expression following host cell entry. *S*. Tm inocula were grown in Luria Bertani broth (LB) with 0.3M NaCl (Sigma-Aldrich) at 37°C for 12h overnight, in the presence of 50µg/ml ampicillin in cases of plasmid-carrying strains. A 1:20 dilution was subcultured in LB/0.3M NaCl without antibiotics at 37°C for 4h. For infection of IEC monolayer cultures and live imaging, the 4h subculture was diluted to 1.0x10^8^ CFU per ml in DMEM/F12 without antibiotics, of which 10µl was used for each infection, resulting in 1x10^6^ CFUs per monolayer. For infection of IEC monolayers in the 96-well plate format, the 4h subculture was diluted to 2x10^8^ CFU/ml in DMEM/F12 without antibiotics, of which 50µl was used for each infection, resulting in 1x10^7^ CFUs per well.

### Live-cell imaging of IEC monolayer infections

Live-cell imaging was performed on a custom-built upright microscope, based on the Thorlabs Cerna upright microscopy system (Thorlabs Inc., Newton, NJ, USA), with a heated 60/1.0 NA Nikon CFI APO NIR objective (2.8mm WD) and a Nikon d-CUO DIC oil condenser (1.4 NA) controlled by Micro-Manager 2.0 (75). Images were acquired with an ORCA-Fusion camera (C14440-20UP; Hamamatsu Photonics, Hamamatsu City, Japan), with a final pixel size of 109nm. Transmitted light was supplied by a 530nm Thorlabs LED (M530L3) to minimize phototoxicity and chromatic aberrations. The imaging chamber was maintained at 37°C with moisturized 5% CO2 air passing over, and using an objective heater. IEC monolayers grown within AICs were placed in 35mm glass-bottom dishes (#3220.110.022, Cellvis, Mountain View, CA, USA) in 3ml DMEM/F12 without antibiotics in the microscope’s light path and allowed to equilibrate for 30min. The *S*.Tm inoculum was then added directly underneath the objective, and imaging was started immediately, with 20ms exposure times and 15sec frame rates for differential interference contrast imaging. The resulting time-lapse movies were analyzed manually using Trackmate for ImageJ (76) to track the time of attachment, division, invasion and/or detachment for individual bacteria. The track information was exported and used to generate dendrogram plots using custom R scripts based on the ‘data.tree’, ‘dendextend’, and ‘stats’ packages.

### Fixed IEC monolayer fluorescence imaging

IEC monolayers grown in AICs were fixed in 4% paraformaldehyde for 30min and permeabilized with 0.1% Triton X-100 for 10min. The monolayers were stained with phalloidin-AlexaFluor488 (Thermo Fisher) and DAPI (#D9542, Sigma-Aldrich) for 30min in PBS. Subsequently, Anodisc alumina membranes were extracted from the AIC membrane holders, washed, and mounted under a 0.17µm coverslip in Mowiol 4-88 (Sigma-Aldrich). For imaging of the effector mutant infections, IEC monolayers grown on PET transwell inserts were infected with 1x10^6^ constitutive *mCherry* expressing *S.*Tm *wt*, *ΔinvG*, or *ΔsipA* bacteria. 60min p.i., the monolayers were washed, fixed in 4% paraformaldehyde for 30min, and stained with Salmonella O Antiserum Factor 5 (1:200, Difco/Miclev), goat-anti-rabbit-AF488 (1:400, Thermo Fisher), DAPI, and phalloidin-AlexaFluor647 (Thermo Fisher). For imaging the PET membranes were removed from their support with a scalpel and mounted under a 0.17µm coverslip in Mowiol 4-88. For lectin stainings, IEC monolayers grown on PET transwell inserts were fixed in 4% paraformaldehyde for 30min and subsequently washed in PBS. The monolayers were stained with either WGA-AlexaFluor647 (#11510826, Thermo Fisher/Invitrogen), UEA-I-FITC (#L9006, Sigma-Aldrich), or SNA-I-Cy3 (#CL-1303-1, Vector Laboratories), combined with DAPI, for 1h in PBS. The monolayer membranes were washed, removed from the plastic transwell holder, and mounted under a 0.17mm coverslip in Mowiol 4-88. The samples were imaged on a Nikon Eclipse-Ti2 microscope equipped with a 100x Plan Apo oil objective (1.45 NA; 110nm final pixel size; Nikon), and a back-lit sCMOS camera (Prime 95B, Photometrics) mounted to a spinning disk module (X-ligth V2, Crest optics). Bright-field images were collected using differential interference contrast (DIC), while fluorescence was excited using a light engine (Spectra-X, Lumencor). Filter set 89402m was used for DAPI and Alexa647, ET600/50m for Cy3, and ET525/50m for FITC (all from Chroma Thechnology Corp). For measurements of nucleus-to-apical plasma membrane distances, spinning disk confocal z-stacks with a 0.3μm step size were acquired for fixed IEC monolayer samples stained for WGA and DAPI, using a 100x oil objective. The z-distance between maximum DAPI signal (nucleus) and maximum WGA signal (apical surface) was measured in Fiji. For maximum-resolution confocal microscopy, samples were imaged on a Zeiss LSM700 inverted point-scanning microscope with a 63x/1.4 NA Plan Apo oil immersion objective using a voxel size of 70.6nm (x and y) by 0.34µm (z-sections). Images were processed and fluorescence intensities quantified using Fiji. For the effector mutant intracellular/extracellular analysis, 7-plane z-stacks were taken in a 4x4 xy-grid with a voxel size of 108nm (x and y) by 2.0µm (z-sections). Fluorescent images were then analyzed in Fiji with a custom script to quantify the mCherry and AlexaFluor488 (AF488) positive spots. The resulting data was analyzed using R to quantify the number of intracellular (mCherry+AF488-) and extracellular (mCherry+AF488+) bacteria. Since mCherry expression efficiency differs somewhat between the effector mutant strains, the counted number of mCherry+ bacteria was corrected by dividing the count by the mCherry expression ratio from the extracellular bacteria (number of mcherry+AF488+ / total number of AF488+). Finally, the number of bacteria was normalized to the total number of bacteria counted per strain in all conditions within one donor-replicate to account for differences in cell and bacterium numbers.

### Fixed IEC monolayer SEM imaging

IEC monolayers grown in the indicated media within AICs were left untreated, or infected with *S*.Tm *wt* for 40min. To fix the samples, monolayers were gently washed once with prewarmed PBS and fixed at 4°C overnight with 2.5% glutaraldehyde (Sigma Aldrich) in 0.1M PHEM buffer [60mM piperazine-N,N’-bis(2-ethanesulfonic acid), 25mM HEPES, 10mM EGTA, and 4mM MgSO .7H 0, pH 6.9]. Prior to SEM imaging, the Anodisc membranes with the fixed monolayers were dehydrated in a series of graded ethanol, critical point dried (Leica EM CPD300), and coated with 5nm platinum (Quorum Q150T-ES sputter coater). The sample morphology was examined by a field emission scanning electron microscope (FESEM; Carl Zeiss Merlin) using in-lens and in-chamber secondary electron detectors at an accelerating voltage of 4 kV and a probe current of 100 pA.

### Gentamicin protection assays and CFU plating

Gentamicin protection assays were carried out using completely confluent cell monolayers cultured on 0.32cm^2^ surface areas. For infection of HeLa cells, 2x10^4^ cells were seeded in 96-well plates 24h prior to infection. For infection of IEC monolayers, 2.5x10^5^ cells were seeded atop PET transwell inserts and grown as specified above in either OGM medium for 3 days, or ENR medium for 7 days. Infections were carried out adding a 10^7^ CFU *S*.Tm wt inoculum (spiked with a non-invasive *ΔinvG* strain to ensure specificity). Following 40min of infection, the monolayers were washed with fresh culture medium, incubated with culture medium containing 200μg/ml gentamicin for 40 min, washed with PBS, lysed in 0.1% sodium deoxycholate (Sigma-Aldrich), and the lysates were serially diluted and plated on LB agar containing appropriate antibiotics.

### IEC monolayer infection assay in 96-well format

IEC monolayers grown in 96-well plates in the indicated medium where washed and incubated in DMEM/F12 without antibiotics for 2h prior to infection. The *S*.Tm *wt*/p*ssaG*-GFP inoculum, prepared as described above, was added directly to the wells followed by incubation for 20min at 37⁰C with 5% CO_2_. After infection the monolayers were washed and incubated in DMEM/F12 supplemented with 100µg/ml Gentamicin for 3h at 37⁰C with 5% CO_2_. Infected monolayers (and uninfected controls) were fixed in 2% paraformaldehyde for 30min, permeabilized with 0.1% Triton X-100, washed and stained with phalloidin-AlexaFluor647 and DAPI for 30min in PBS. After staining, the monolayers were washed with PBS and imaged using the Nikon Ti2 microscope described above, but through a 40X/0.6 NA Plan Apo air objective (pixel size 0.29µm). Infection foci quantification was performed using Cellprofiler (77). For the StcE treatment assays, ENR-grown IEC monolayers were preincubated in 10µg/mL StcE (active) or StcE E447D (inactive) in DMEM/F12 for 2 hours at 37⁰C with 5% CO_2_, prior to infection.

### Expression and purification of StcE and inactive StcE E447D

Tuner (DE3) competent cells (70623, Sigma-Aldrich) were transformed with pET28b-StcE-Δ35-NHis or pET28b-StcE-E447D-Δ35-NHis (kind gift from Prof. Carolyn Bertozzi; (56)) and grown on LA agar with kanamycin at 37°C overnight. A single colony was pre-cultured in 10ml LB/Kanamycin overnight, and the preculture expanded in 1L LB/Kanamycin until an OD_600_ of 0.85 was reached. Protein production was induced with 0.2mM IPTG at 30°C overnight. Bacterial cells were centrifuged at 3500 x g for 20min at 4°C, resuspended in 20ml ice-cold PBS, and centrifuged at 3500 x g for 20min at 4°C. Cell pellets were resuspended in 20ml ice-cold Binding Buffer (20mM sodium phosphate, 300mM NaCl, 20mM Imidazole, pH 7.4) containing 2.5X Roche Complete EDTA-free protease inhibitor cocktail (#11873580001, Sigma-Aldrich). The bacterial slurry was sonicated for 8 x 30sec at 50% duty in a water bath maintained at 4°C. Lysates were centrifuged at 22000 x g at 4°C for 20min and poured over 4ml of HisPur Cobolt resin (#89964, Thermo Fisher Scientific). The slurry was rotated at 4°C for 1h and spun down at 700 x g for 2min. The resin was washed three times with Binding Buffer including the protease inhibitor cocktail, and the bound protein was eluted at 4°C with three subsequent 15min elutions using 3ml of Elution Buffer (20mM sodium phosphate, 300mM NaCl, 500mM Imidazole, pH 7.4). Elution fractions were pooled and dialyzed against 5L PBS at 4°C overnight, followed by a second round of dialysis against 5L PBS for 4h at 4°C.

### Barcoded consortium infections of apical-out 3D enteroids in suspension

Medium sized human enteroids were dislodged from the matrigel domes and incubated in Cell Recovery Solution (Corning, #354253) for 1h on ice on a rotating table. After allowing enteroid sedimentation by gravity, the supernatant was removed and the pellet washed with DMEM-F12/0.25% BSA. The pellet was then gently re-suspended in OGM or ENR media without Matrigel to promote eversion (apical-out; (32)) and enteroids were aliquoted in ultra-low attachment 24-well tissue culture plates (Corning Costar; #CLS3473-24EA). Apical-out 3D suspension cultures were incubated at 37°C with 5% CO_2_ for 3-4 days (for OGM condition), or 6-7 days (for ENR condition) prior to infection. Shortly before the infection, apical-out enteroids were washed with DMEM-F12/0.25% BSA, using 25µm mini cell strainers (Funakoshi, #HT-AMS-14002). Once recovered from the strainers, enteroids were resuspended in DMEM/F12/0.25%BSA and aliquoted in ultra-low attachment 24-well tissue culture plates (Corning Costar; #CLS3473-24EA). The indicated tagged *S*.Tm strains were mixed in equal ratio and diluted to obtain a total MOI of 40. Bacteria were added to each well and incubated for 20min. Subsequently, enteroids were washed 4 times with DMEM-F12/0.25% BSA using 25µm mini cell strainers and incubated with media containing 200µg/ml gentamicin in the lids of 14 ml falcon tubes for 1h. Infected enteroids were washed 6 times with DMEM-F12/0.25% BSA, recovered from the strainers and lysed in 0.1% Na-deoxycholate by homogenization with a Tissue Lyser (Qiagen). The recovered intracellular bacterial populations were enriched ON at 37°C in 2ml LB. A diluted culture of the inoculum consortium was also enriched in the same condition and used as the input reference. Genomic DNA from enriched cultures was extracted using the GenElute Bacterial Genomic DNA kit (Sigma; #NA2110-1KT). For tag quantification, qPCR was performed using Maxima SYBR green/ROX qPCR master mix (2X) (Thermo Fisher Scientific, #K0222) on a CFX384 Touch Real-Time PCR Detection System (Biorad), using tag-specific primers (20). The relative abundance of each strain was normalized to the abundance in the inoculum and expressed as % of total intracellular population. Infection indexes were calculated by the formula (1-(mean relative abundance*^wt^*/mean relative abundance*^mutant^*)) in relation to the mean of the *S*.Tm*^wt^*strains (value 0).

### Statistics

Statistical significance testing employed the two-sided t-test with Holm-Bonferroni post-hoc, the Kruskal-Wallis test followed by Dunn’s post-hoc with Benjamini-Hochberg or Holm-Bonferroni correction, or the unpaired Wilcoxon rank-sum test. For barcoded consortium infections, significance testing employed a paired t-test applied to the relative input and output abundances of each strain, normalized by the respective mean *wt* strain abundance. The statistical tests are indicated in the respective figure legends along with p-value specifications.

## Author contributions

Conceptualization: JMvR, ACCL, MLDM, MES; Methodology: JMvR, ACCL, MC, JE, AB, MLDM; Investigation: JMvR, ACCL, MC, AB, AN, MLDM; Formal analysis: JMvR, ACCL, MC, AN, MLDM; Interpretation: JMvR, ACCL, MC, PA, TP, MLDM, MES; Resources: RH, MSu, MSk, WG, DLW, PMH, PA, TP, MES; Project administration: ACCL, RH, MSu, MSk, WG, DLW, PMH, MLDM, MES; Supervision: PA, MLDM, MES; Funding acquisition: PA, MES; Visualization: JMvR, ACCL, MC, MLDM; Writing – original draft: JMvR, ACCL, MES; Writing – reviewing and editing: all authors.

## Acknowledgments

The authors thank members of the Sellin laboratory for fruitful discussions and the staff at the Uppsala University Hospital surgical units for assistance with intestinal tissue sample collection. We are grateful to the BioVis platform at Uppsala University for support with confocal fluorescence microscopy and the Umeå Centre for Electron Microscopy (UCEM) for support with SEM imaging. Transcriptomic analyses were conducted at the National Genomics Infrastructure (NGI) Sequencing Facility, SciLifeLab, Uppsala. This study was funded by grants from the Swedish Research Council (2017-01951 and 2020-01586 to PA, 2018-02223 and 2022-01590 to MES) and the Swedish Foundation for Strategic Research (FFL18-0165 to MES). AB, PA and MES also acknowledges financial support from Uppsala Antibiotic Center (UAC PhD grant 2021) and MC acknowledges support by a PhD scholarship from the Ministry of National Education of Turkish Republic. The funders had no role in design or execution of the research.

## Supplemental figure legends

**Figure S1.**
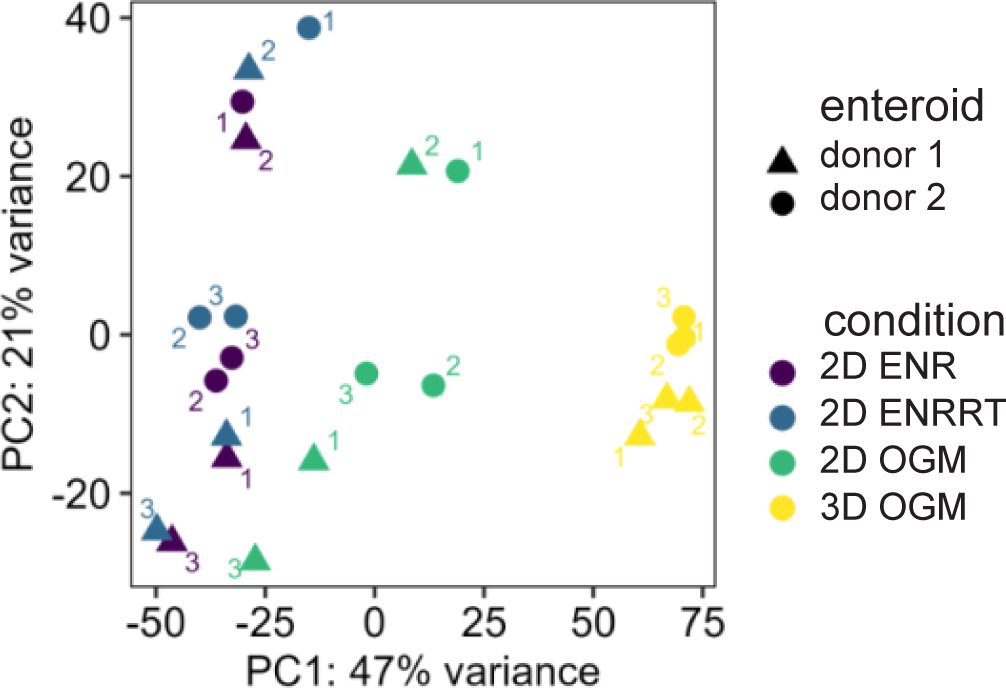
Principal component analysis of transcriptomes from intestinal epithelial cell layers grown under distinct conditions. Enteroid-derived monolayers from two donors were cultured in either OGM, ENR, or ENRRT and the transcriptomes analyzed at day 7 post-seeding. The corresponding 3D enteroid cultures kept in OGM within Matrigel domes were included as an additional reference sample. Each data point represents one out of three biological replicate samples for each sample group. The donor enteroid line used is indicated by the symbol shape. The principal component analysis shows the separation of 3D OGM, 2D OGM, and 2D ENR/2D ENRRT sample groups along the PC1 axis.

**Figure S2.**
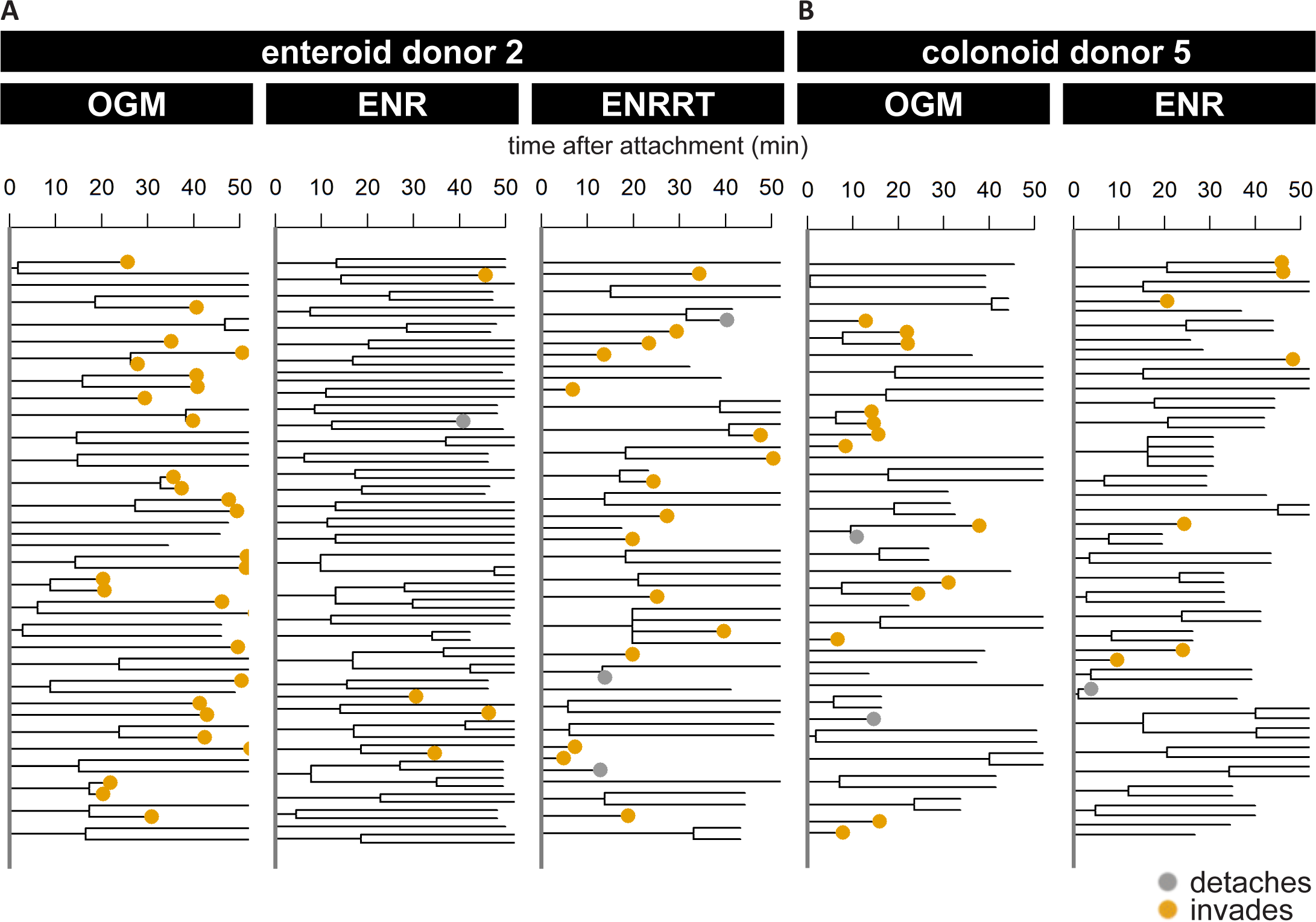
The reduced permissiveness to *Salmonella* invasion upon intestinal epithelial cell layer maturation is observed across different donor cultures. Quantification of the fates of individual *S*.Tm after initial attachment to the epithelial surface. Data derive from time-lapse imaging of **(A)** enteroid-(donor 2) and **(B)** colonoid-derived (donor 5) IEC monolayers, grown in OGM, ENR or ENRRT within apical imaging chambers and infected with *S*.Tm *wt* at MOI 3 for 1h. The behavior of individual surface-bound *S*.Tm in a representative time-lapse movie from each condition is visualized as dendrograms. The dendrograms indicate the timing of bacterial division (track split), detachment (grey dot), invasion (yellow dot), and remaining bacteria at the end of the movie or loss of focus (bare branch). Also for these donor cultures, IEC maturation (ENR condition) confer markedly reduced number of *S*.Tm invasion events.

**Figure S3.**
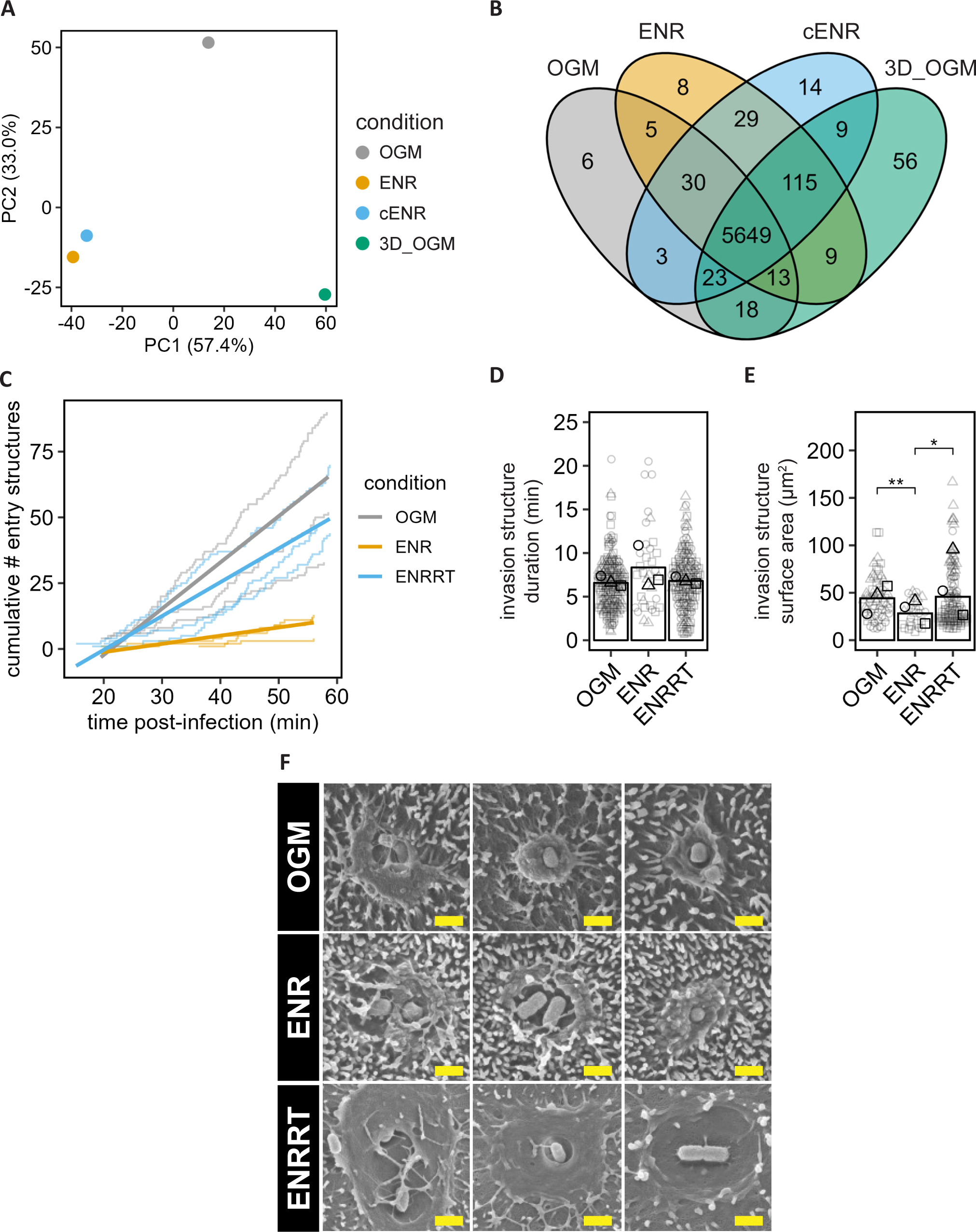
Proteome characteristics and permissiveness to *Salmonella* invasion across intestinal epithelial cell layers grown under distinct conditions. **(A-B)** Enteroid-derived 2D monolayers were cultured in either OGM, ENR, or cENR and the global proteomes analyzed. A corresponding 3D enteroid culture kept in OGM within Matrigel domes was included as an additional reference sample (3D_OGM). **(A)** Principal component analysis of global proteome samples. Each data point represents the average of three biological replicate samples for each condition, with all samples generated from the same donor line. The principal component analysis, based on all quantified proteins, shows the separation of 3D OGM, 2D OGM, and 2D ENR/cENR samples along the PC1 axis. **(B)** Venn diagram showing the overlap among all quantified proteins between the sample groups. **(C-E)** Time-lapse imaging of enteroid-derived IEC monolayers grown in OGM, ENR, or ENRRT within AICs and infected with *S*.Tm *wt* at MOI 3 for 1h. The panels show quantification of **(C)** the cumulative number of entry structures detected over time, **(D)** entry structure duration, and **(E)** maximal entry structure area. In D-E, each symbol represents one entry structure pooled from 3 replicate time-lapse movies for each condition. Statistical analysis; Kruskal-Wallis test followed by Dunn’s post-hoc with Holm-Bonferroni correction. *p-value = 0.0159, **p-value = 0.00819. **(F)** Representative scanning electron microscopy images of *S*.Tm-elicited apical entry structures in OGM, ENR, and ENRRT-grown monolayers. Scale bars, 1µm. Note that for panels C-F, the corresponding data for OGM and ENR-grown monolayers have been replotted from Fig 3 for reference. The results show that IEC monolayers kept in an immature state (OGM), or differentiated towards an M-cell-like phenotype (ENRRT), are more permissive to *S*.Tm invasion than monolayers differentiated towards an enterocyte phenotype (ENR).

**Figure S4.**
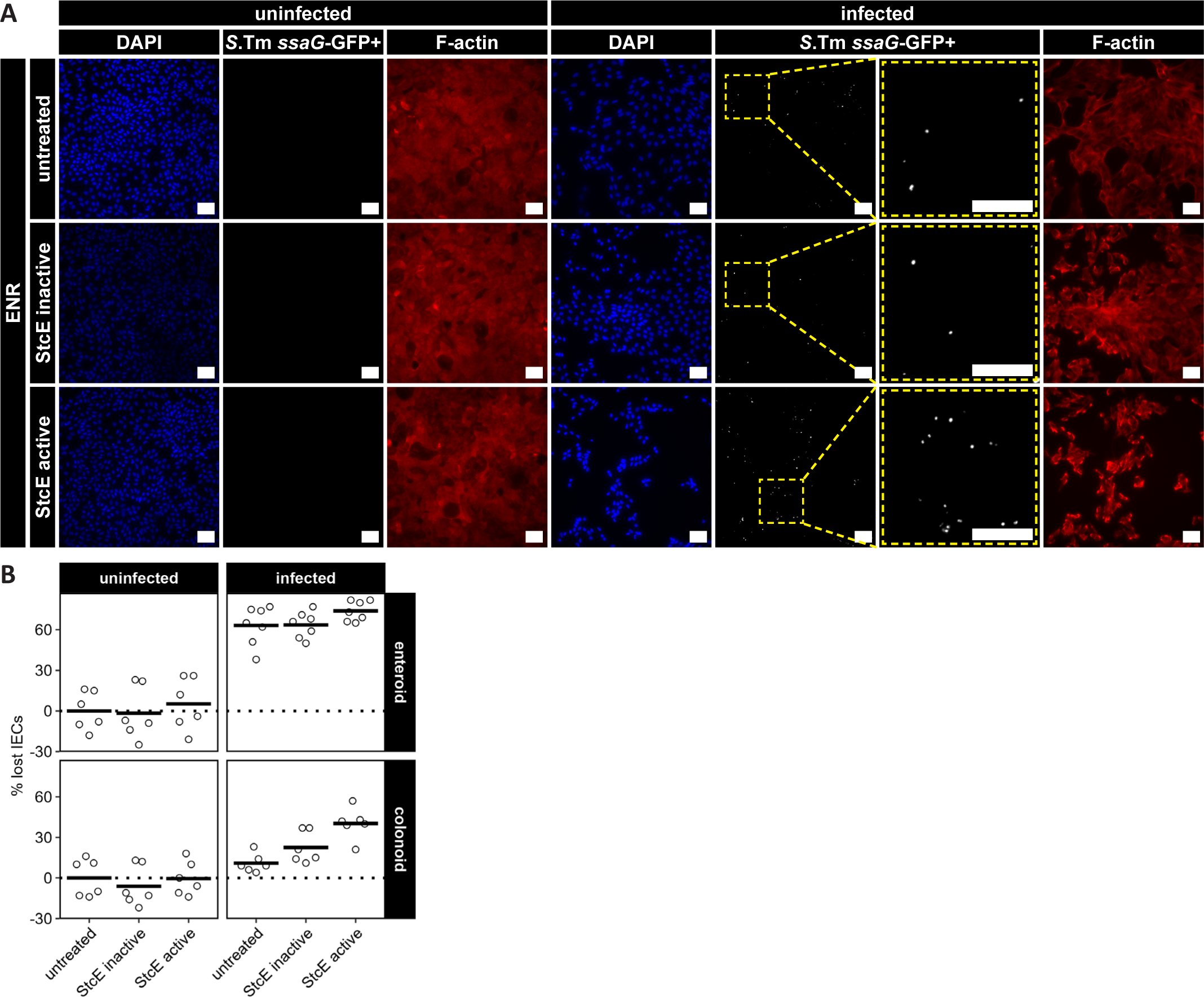
Morphology and cell death in intestinal epithelial cell layers pretreated with the StcE enzyme prior to *Salmonella* infection. Enteroid-and colonoid-derived IEC monolayers were grown in ENR within microwells, left untreated, treated for 2h with enzymatically inactive StcE, or active StcE, and subsequently infected with the *S*.Tm *wt*/p*ssaG*-GFP reporter strain at MOI ∼40 for 20min, followed by 3h further incubation for reporter maturation. **(A)** Representative fluorescence micrographs of monolayers stained with DAPI (nuclei) and phalloidin (F-actin). Second last column shows blow-ups of boxed regions. Scale bars, 50µm. **(B)** Quantification of the percentage of IECs lost by the time-point of analysis. *S*.Tm infection results in significant death and loss of infected IECs within this experimental window.

